# Wild pigs mediate far-reaching agricultural impacts on tropical forest soil microbial communities

**DOI:** 10.1101/2021.09.10.459828

**Authors:** Francis Q. Brearley, Hokyung Song, Binu M. Tripathi, Ke Dong, Noraziah Mohamad Zin, Abdul Rahim Abdul Rachman, Kalan Ickes, Jonathan M. Adams, Matthew S. Luskin

**Affiliations:** Department of Natural Sciences, Manchester Metropolitan University, Chester Street, Manchester, M1 5GD, UK; Department of Earth and Environmental Sciences, The University of Manchester, Manchester, M13 9PL, UK; Department of Biological Sciences, Seoul National University, Gwanak-ro1, Seoul, 08826, South Korea; Korea Polar Research Institute, Incheon, 21990, South Korea; Department of Life Sciences, Kyonggi University, Suwon, 16227, South Korea; Programme of Biomedical Science, Faculty of Health Sciences, Universiti Kebangsaan Malaysia, Kuala Lumpur, Malaysia; Faculty of Science, Technology, Engineering and Mathematics, International University Malaya-Wales, Jalan Tun Ismail, 50480, Kuala Lumpur, Malaysia; Central, South Carolina, USA; School of Geography and Oceanography, Nanjing University, Nanjing, China; Forest Global Earth Observatory, Smithsonian Tropical Research Institute, Washington, DC, USA; Asian School of the Environment, Nanyang Technological University, Singapore, Singapore; School of Biological Sciences, The University of Queensland, St Lucia, Brisbane, QLD 4072, Australia

## Abstract

Edge effects, the altered abiotic and biotic conditions on the borders of natural areas, rarely extend more than a few hundred meters. Edge effects have rarely been linked to altered soil biota, which shape ecosystem processes including carbon storage, biogeochemical cycling, and plant performance. Here, we investigated if agriculturally-mediated increased wildlife populations affect soil biotic communities at a distance well over that of estimated edge effects when they move between agriculture and natural habitats using a 22-year fenced exclusion experiment in a primary rainforest in Peninsular Malaysia. We found that the presence of wildlife (mainly native pigs (*Sus scrofa*) that crop-raid in nearby oil palm plantations) was associated with higher bacterial diversity, and an altered community composition (mediated by changes in soil pH), and reduced abundances of symbiotic ectomycorrhizal fungi compared to soil in exclosures. There were only minor effects of pigs on soil chemistry or microclimate, so we suggest that changes in soil communities are driven by pigs’ leaf litter removal and alterations to plant composition. Our study highlights that indirect effects from agriculture can be transferred by wildlife >1 km into protected areas and this could have important repercussions for ecosystem processes and plant-soil feedbacks.

## INTRODUCTION

Agricultural encroachment into forested areas is a pervasive global phenomenon that has a clear and direct impact on above-(1,2) and below-ground (3) biota and ecosystem processes, especially in tropical forests. It is more challenging to assess how agricultural expansion indirectly affects natural areas over larger spatial scales, such as the cryptic degradation from edge effects. One example is cross-boundary ecological cascades, wherein adjacent ecosystems – first appearing to be distinct – are actually linked through the transport of nutrients (e.g. via floods or mobile animals) or interactions with wildlife that moves across ecotones (4). With over 70 % of remaining forests now lying within 1 km of an edge (5), there is an urgent need to understand how edge effects reshape the linkages between above- and below-ground biota and the scale at which they operate.

Wildlife responses to edges are related to their unique habitat preferences, the local hunting intensity and preferences, and some crop-raiding wildlife can even benefit from supplemental foraging in nearby farmlands (4,6-9). Edge effects also produce a range of impacts on soils that are often mediated by microclimate and light (10), as well as plant species composition (e.g. 11-13). However, these edge effects have rarely been documented beyond a few hundred meters (10,14,15). Wildlife plays an important role moderating nutrients like nitrogen and phosphorus, and soil biota through deposition of excrement and carcases (16-19). Wildlife also affects soil physical environments through biopedoturbation (20-22) and plant-soil interactions via herbivory or nest building (23-26).

Here, we investigate the potential for wide-ranging wildlife to transfer far-reaching edge effects on soil microbial communities in distant ‘interior’ primary forests at distances >1 km from the nearest edges. We conducted our study in a primary Malaysian rainforest where native forest-dwelling wild boars (*Sus scrofa*) that forage in nearby oil palm plantations (*Elaeis guineensis*) have elevated densities and are known to disturb nearby forest soils and plant communities (4,27,28). Wild boars (hereafter ‘pigs’) are a key example of a broadly distributed generalist vertebrate that is adaptable to human environments (e.g. forest edges) and strongly affected by humans via hunting (negative) or crop-raiding (positive). Pigs are considered an ‘ecosystem engineer’ due to the major physical soil disturbances via rooting, grubbing (predating larger soil invertebrates), wallowing, trampling and soil compaction (29-30).

We focus on pigs’ influence on microbial community composition and functioning, which remains largely unknown and has myriad links to biogeochemical processes that in turn shape ecosystem properties including carbon dynamics (31). Using a long-term exclosure experiment, we examined three hypotheses regarding the indirect impacts from oil palm-fed pigs on the forest soil microbial communities based on the ecology of pigs and known relationships between soil disturbances and microbial communities:

1. First, we predicted that pig-exposed soils would have greater nutrient concentrations due the deposition of urine and faeces and that, together with disturbances caused by removal of understorey plants and leaf litter by pigs, these would be key drivers of microbial community structure in pig-exposed soils.
2. Second, we predicted that soil bacteria would be more impacted than fungi due to changes in nitrogen deposition from excrement influencing bacteria involved in the nitrogen cycle, and removal of understorey plants and leaf litter, which increases light penetration and likely has a drying effect on soils to which bacteria are more sensitive than fungi (32).
3. Third, we predicted that whilst fungi would be less influences by the presence of pigs than bacteria, symbiotic ectomycorrhizal (EcM) fungi would be reduced in pig-exposed soils because pigs preferentially remove dipterocarp seedlings (23) that are associated with these fungi.

## RESULTS

There were low concentrations of all soil nutrients measured, which is typical for Southeast Asian rain forests on similar substrates (Table S1). There were no significant differences between the exclosures and the open-control areas with the exception of soil pH that was 0.15 pH units more acidic within the exclosures.

Bacterial richness was 13 % lower in exclosure soils (t-test, P = 0.042; Fig 1a). The relative abundance of Acidobacteria increased (t test, P = 0.013; Fig 2a) but there was lower relative abundance of Proteobacteria, Actinobacteria, Planctomycetes and Gemmatimonadetes (all P < 0.05; Fig. 2). At the subphylum level, we also observed significant differences in the relative abundance of dominant bacterial taxa, for example, the relative abundance of bacterial families Solibacteraceae and Rhabdochlamydiaceae increased, whilst abundance of other families such as Bradyrhizobiaceae and Xanthomonadaceae, including members with biological N_2_-fixation capacities, declined in exclosure soils (Table S2). The bacterial community composition differed between the exclosures and the open-controls (ANOSIM R = 0.216, p 0.009; Figs 2 & 3a) and was influenced by soil pH (Fig. 3a). Soil carbon and potassium also influenced the bacterial community composition, but these did not differ between the exclosure and open-control soils (Table S1). There was no clear influence of the exclosures on bacterial community functioning as measured by predicted gene abundance (ANOSIM R = 0.081, p = 0.23; Fig 3c).

**Figure 1:**
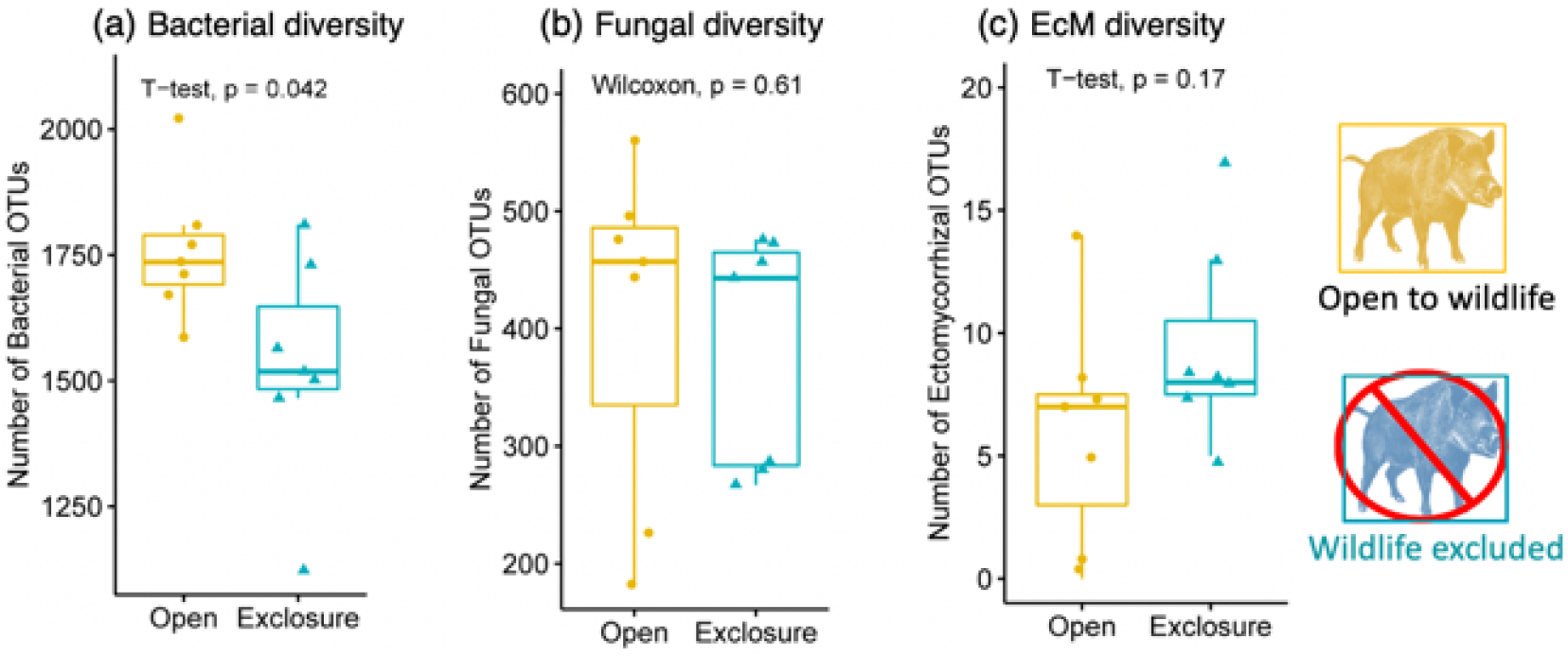
The long-term effects of wildlife (primarily pigs) on soil microbial diversity at Pasoh Forest Reserve in Peninsular Malaysia. Samples are separated by whether they were taken from open-control plots where there were many pigs (yellow dots) versus within fenced exclosures without pigs (blue triangles). EcM = Ectomycorrhizal fungi.

**Figure 2:**
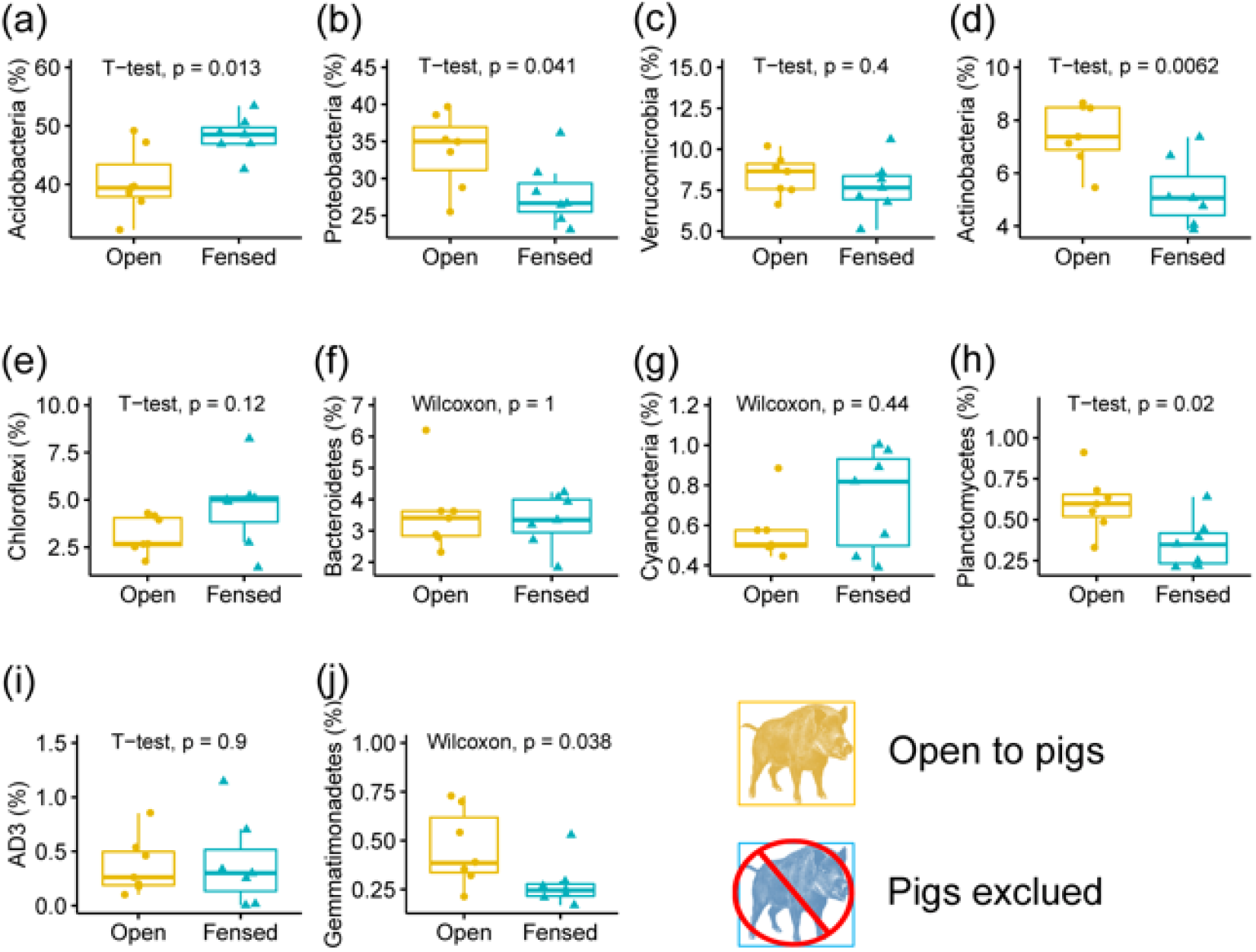
Influence of wildlife (primarily pigs) on the relative abundance of soil bacterial phyla at Pasoh Forest Reserve in Peninsular Malaysia (interpretation is the same as Fig. 1).

**Figure 3:**
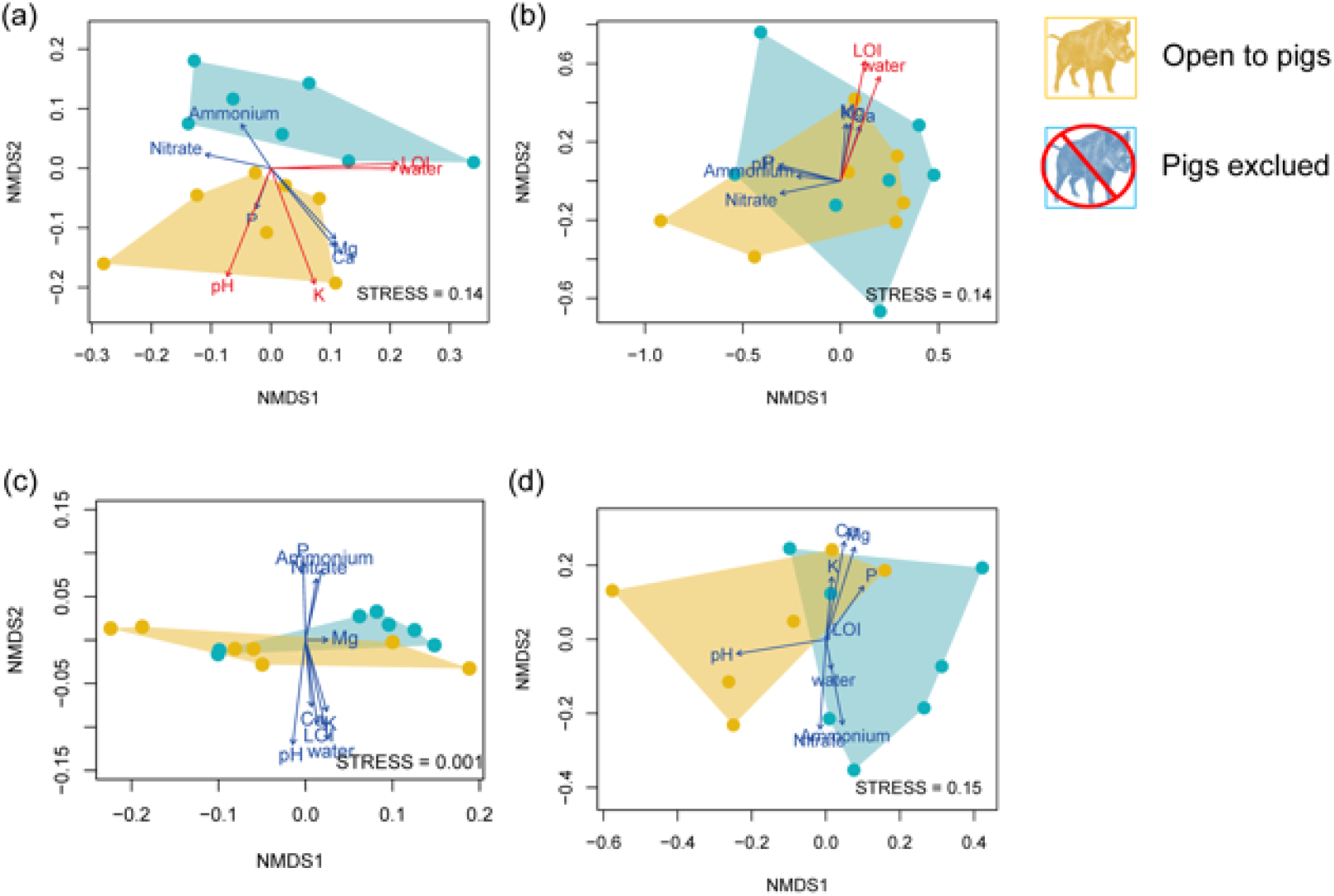
NMDS ordinations showing the influence of wildlife (primarily pigs) on the soil microbial community composition and function at Pasoh Forest Reserve in Peninsular Malaysia. **(a)** Bacterial taxa **(b)** Fungal taxa **(c)** Bacterial gene abundance **(d)** Ectomycorrhizal fungal taxa. Samples are separated by whether they were taken from open-control plots where there were many pigs (yellow dots) versus within fenced exclosures without pigs (blue triangles). Environmental factors that have a significant influence are marked in red. LOI = loss-on-ignition (%).

The fungal community did not differ significantly between the exclosure and open-control soils in terms of diversity (Wilcoxon test, P= 0.61; Fig 1b), phylum abundance (Wilcoxon test, all P > 0.05; Fig S1), community composition (ANOSIM: R = -0.043, p = 0.70; Fig 3b) or guild composition (Fig. S2). However, the ectomycorrhizal (EcM) fungal community composition differed between wildlife treatments (ANOSIM p = 0.03; Fig 3d), becoming 1.3 times more diverse (t-test, P= 0.17; Fig. 1c) and 2.9 times more abundant in exclosure soils (Wilcoxon test, P= 0.16; Fig. S2b). In particular, although the abundance of the most common EcM family Russulaceae was variable it was, overall, 2.5 times more abundant in the exclosure soils (Wilcoxon test, P= 0.38). The fungal community composition (including all species) was influenced by soil carbon (Fig. 3b) but none of the soil chemical variables significantly influenced the EcM community structure.

## DISCUSSION

Our study is the first to link agricultural incursions to altered soil microbial community composition in adjacent habitats at distances >1 km from edges, one of the furthest soil-related edge effects yet recorded. These far-reaching edge effects were mediated by crop-raiding native pigs that fed on oil palm fruits in adjacent plantations and then return to forests. The biotically-driven impacts by pigs that we report are distinct from more well-documented edge effects related to habitat and abiotic conditions, since all our sampling locations were equidistant from edges and microclimatic differences were not detected greater than 100 m from the edge at our site (33). Instead, we posit that pigs impact soil microbial communities by disturbing soils, leaf litter, and understory vegetation and altering the plant community composition (4,23,34). For example, previous work at our site has shown pigs reduce trees with symbiotic root-associated ectomycorrhizal (EcM) fungi (e.g. Dipterocarpaceae) and facilitate lianas that are rarely reported to have EcM associations (35). As predicted, we found wildlife exclusion was associated with altered EcM fungi communities and a greater relative abundance of EcM fungi. Wildlife exclusion was also associated with reduced soil bacterial diversity which led to a greater community change than for the fungal community. Prior work has found invasive pigs reduced soil bacterial diversity in Hawaii (36) – the opposite trend we observed from native pigs - and, in New Zealand, invasive pig grubbing may increase the relative abundance of fungi over bacteria (37) which our results supported.

We did not detect higher soil nutrient concentrations due to pig excrement (urine and faeces) deposition in open-control plots as we had predicted. Such equivocal results align with work finding a wide variety of impacts from wildlife on soil nutrients (17,38,39). It is also possible that additional nutrients from wildlife excrement could have been balanced with loss from leaf litter removal. Soil pH was slightly (0.15 pH units) more acidic in exclosure soils but - as has previously been noted by others (40,41) - this was associated with altered bacterial community composition. As predicted, we found varying trends in relative abundance of bacterial taxa, for example, the family Solibacteraceae (phylum Acidobacteria) dominated in exclosure soils, while the family Vicinamibacteraceae (phylum Acidobacteria) was abundant in open-control soils. The relative abundance of Solibacteraceae is reported to decline with increasing soil pH (42) and organic fertilization (e.g. manure) (43), while that of Vicinamibacteraceae is positively correlated with pH (42) and had a higher prevalence in nutrient-amended soils (44). Contrary to our prediction, the relative abundance of bacterial families such as Bradyrhizobiaceae and Xanthomonadaceae, which include members with biological N_2_-fixation capacities, increased in open-control soils. However, N_2_-fixing members of these families are also known for their denitrifying abilities (45), which suggest that nitrogen deposition from urine and faeces from the pigs could have increased their abundance together with other denitrifying bacterial taxa such as in Rhodospirillaceae (46). Furthermore, in accordance with our prediction, we found a dominance of drought-tolerant Actinobacterial taxa (47,48) in open-control soils.

We had predicted soil nutrients, water, and soil carbon, which are associated with the energy available for micro-organisms, would drive differences in soil microbial communities. However, while we found that bacterial community composition was associated with soil carbon and potassium, these attributes did not differ between exclosures and open-control soils suggesting these factors did not explain differences in bacterial communities between exclosure treatments. Instead, bacterial community change was more likely associated with abiotic conditions, including leaf litter removal and increased light penetration from a browsed and trampled understory vegetation that may cause increased light penetration and soil drying.

In support of our third hypothesis, the EcM fungal community differed significantly between treatments, and open-control soils were associated with lower relative abundance and diversity of EcM fungi, although there was borderline statistical significance due to only having seven replicates and the potential to improve fungal guild assignment via FUNGuild (49). Pigs may directly consume some fungi or, more likely, the disproportionate removal of trees with symbiotic relationships with EcM fungi (23) is key here. The decline in EcM fungi may cause plant-soil feedbacks that reduce regeneration of EcM-dependent plant species such as dipterocarps (50) and therefore influence future patterns of forest composition. Future work testing if EcM fungi and their plant symbionts differ in unhunted forests with abundant pigs (e.g. Pasoh) compared to hunted forests were pigs are rare, such as in Lambir Hills in Malaysian Borneo (51) would be of great interest.

In summary, our study is notable for documenting that cryptic biotically-driven edge effects are mediated by wide-ranging wildlife and affect soil microbial community composition. The magnitude of impacts we observed at our site is linked to elevated pig populations associated with oil palm plantations and low hunting pressure(4), thereby creating a cross-boundary ecological cascade from agriculture to pig to soils that extends > 1 km. Pigs are common throughout Asia and invasive globally (29,30,52,53), so our findings may be generalizable beyond Malaysia, and to other mobile crop-raiding wildlife species besides pigs (7,54). The onslaught of African Swine Fever in Asia, which has spread to wild pigs, may reduce pigs’ abundance, and provide opportunities for natural experiments on the ecological impacts of losing pigs (55). Other future research may examine associations between soils and volant animals that are often wider ranging, and examine feedback loops between altered soils and plant composition. We conclude with a warning that far-reaching edge effects may produce consequential changes to ecosystem properties and processes performed by soil microbes, as well as alter plant performance and community composition in the future.

## METHODS

### Study site

The study was conducted at the Pasoh Forest Reserve, Negeri Sembilan, Peninsular Malaysia (2°59’ N, 102°18’ E) where the mean annual precipitation is approximately 1800□mm (56). In the study area, the soils are developed over shale, granite and alluvial parent materials with a generally gentle topography and a fairly homogenous vegetation composition (56). The lowland evergreen rain forest core of the reserve is a 600 ha tract dominated by Dipterocarpaceae and typical of much of the broader region (56). Oil palm plantations surround the reserve on three sides (extending for 2 to 10 km away from the reserve) with the northern side abutting a contiguous area of selectively logged lowland and hill forest. Pasoh supports a diverse wildlife community (4) but pigs (*Sus scrofa*) are by far the most common mammal and present at very high densities of 27-47 per km^2^ (4, 52). Pigs are known to cause soil disturbances in the forest that is tied to their nightly traveling to oil palm plantations to crop-raid (27).

### Wildlife exclosure experiment

Eight open-topped exclosures were constructed in 1996 along the southern edge of the 50-ha permanent Forest Dynamics Plot, and 1.3 km from the nearest forest edge (57). The exclosures were 7 m x 7 m, with 1.5 m tall fences made from 4-cm^2^ chain-link metal and surrounded by barbed wire. Each fenced area was paired with two adjacent open-control areas located at least 1 m outside the fences. At the time of this study, seven remained effective and one was damaged by falling trees and was not surveyed. Exclosures are described in more detail by Ickes et al. (34) and Luskin et al. (57).

### Soil sampling and DNA extraction

Surface soil (0-5 cm depth) samples were collected in July 2018 from the seven exclosures and seven of their paired controls. We took soils from four points at the corners of a 1 × 1 m^2^ grid and composited them for further analysis. We avoided sampling areas in the controls that had been grubbed by pigs, as this would have exposed sub-surface soil that is known to have a different microbial community to the upper horizons. Soils were kept chilled for c. 48 hours before DNA was extracted from 0.25 g of each soil sample using a MoBio PowerSoil kit following the manufacturer’s instructions.

### DNA sequencing

Extracted soil DNA was PCR-amplified in duplicate using the high-fidelity Phusion polymerase. A single round of PCR was done using “fusion primers” (Illumina adaptors + indices + specific regions) targeting the V6-V8 region of 16S rRNA gene of bacteria and the internal transcribed spacer (ITS) 2 region of fungi using the B969F & BA1406R primers of Comeau et al. (58) and ITS86(F) & ITS4(R) primers of Op De Beeck (59) respectively. The PCR products were cleaned and normalized using the high-throughput Charm Biotech Just-a-Plate 96-well Normalization Kit and pooled to make one library that was quantified fluorometrically before sequencing. Sequencing library construction and Illumina MiSeq sequencing (2 × 300 bp) were performed at the Integrated Microbiome Resource, Dalhousie University, Canada (https://imr.bio/index.html).

### Bioinformatics

Forward and reverse sequences were assembled using PANDAseq v.2.8 and further sequence processing was performed following the MiSeq SOP in Mothur v.1.32.1 with chimeric sequences removed using chimera.uchime (60-62). Operational taxonomic units (OTUs) of bacterial 16S rRNA gene sequences were assigned based on the OptiClust algorithm using Mothur v.1.40.5 with a 97 % similarity threshold and OTUs of fungal ITS sequences were assigned based on the UCLUST algorithm using QIIME v.1.9.1 with a 97 % similarity threshold (63,64). Singleton sequences were removed. Bacterial sequences were then classified based on EzBioCloud database v.2018.05 for bacteria (65) and the UNITE database v.7.2 for fungi (66). To infer the bacterial functions from 16S rRNA gene sequences, we used Phylogenetic Investigation of Communities by Reconstruction of Unobserved States (PICRUSt v. 1.1.2 (67). PICRUSt uses extended ancestral-state reconstruction algorithm to generate the composition of gene families for the subset of OTUs present in Greengenes database v. 13.5 (68). The predicted gene families were then classified into Kyoto Encyclopedia of Genes and Genomes (KEGG) orthologues (69). We used FUNGuild v.1.0 for functional guild classification of fungi (49).

### Soil analyses

In the field, c. 2.5 g fresh soil was added to 20 ml of 1 M KCl, shaken and returned to the field laboratory where it was filtered through a 0.2 µm filter after c. 6 hours. It was then diluted 1:4 and analysed on a Dionex ICS 6000 ion chromatograph for available ammonium and nitrate. The moisture content of fresh soil was determined by heating subsamples to 105 °C for 24 h and the remainder was air-dried and ground to pass a 1-mm sieve. Soil pH was measured by adding 2.5 g of soil to 6.25 ml of deionised water; the mixture was then shaken and left to equilibrate for 24 h before measurement with a Sartorius PB-11 pH meter. Total carbon and nitrogen were determined on a Vario EL Cube elemental analyser. Cations (P, K, Ca and Mg) were extracted from 2.5 g sub-samples that were shaken with 25 ml of Mehlich 3 solution for ten minutes before being filtered and analysed on a Thermo iCAP 6300 Duo inductively coupled plasma optical emission spectrometer with correction by determining moisture content of the air-dried soil by heating subsamples as above.

### Statistical analyses

For diversity analysis, bacterial sequences were subsampled into 23,601 reads and fungal sequences were subsampled into 8,994 reads. To compare relative abundance of phyla and diversity between treatments we used t-tests when data were normally distributed or Wilcoxon rank sum tests when data were not normally distributed. We used Bray-Curtis dissimilarity (based on square-root transformed abundances) to visualize differences in the bacterial and fungal communities between treatments and the KEGG Level 3 gene assignments (69). We drew NMDS (non-metric multidimensional scaling) plots using the ‘metaMDS’ function in R package ‘vegan’ (70). Statistical significance between treatments were tested by Analysis of Similarities test (ANOSIM). We assessed if microbial composition was influenced by abiotic conditions by including environmental vectors (covariates) in the NMDS ordinations using the vegan ‘envfit’ function’.

## ACKNOWLEDGEMENTS

We acknowledge the grant provided by Universiti Kebangsaan Malaysia for molecular work (M1-2018-004). We thank the Forest Research Institute of Malaysia (FRIM), the Negeri Sembilan Forestry Department and the Pasoh Research Committee for maintaining the Pasoh Forest and for research permission. We acknowledge ForestGEO of the Smithsonian Tropical Research Institute and FRIM for maintaining and supporting the Pasoh 50-ha forest dynamics plot and assistance funding for the exclosure experiment. We thank XX anonymous reviewers for useful comments on earlier drafts.

## Supplementary Materials

**Table S1:** Soil chemical analysis (mean ± standard error) within and outside exclosures to prevent the influence of wildlife (primarily pigs) on ecological processes at Pasoh Forest Reserve in Peninsular Malaysia.

| Soil Property | Exclosure | Open-Control | t-test |
| --- | --- | --- | --- |
| pH | 4.03 ± 0.02 | 4.18 ± 0.06 | t=2.42, p=0.032 |
| Loss-on-ignition (%) | 5.71 ± 0.75 | 5.29 ± 0.69 | t=0.42, p=0.69 |
| C (%) | 2.18 ± 0.87 | 2.15 ± 0.73 | t=0.05, p=0.96 |
| N (%) | 0.15 ± 0.02 | 0.14 ± 0.02 | t=0.30, p=0.77 |
| Ammonium ($\mu\text{g g}^{-1}$) | 6.08 ± 10.90 | 1.54 ± 1.13 | t=1.09, p=0.30 |
| Nitrate ($\mu\text{g g}^{-1}$) | 2.43 ± 4.70 | 2.63 ± 4.80 | t=0.08, p=0.94 |
| Available P ($\mu\text{g g}^{-1}$) | 9.19 ± 3.90 | 9.58 ± 3.92 | t=0.18, p=0.86 |
| Available K ($\mu\text{g g}^{-1}$) | 56.8 ± 19.9 | 79.7 ± 43.6 | t=1.27, p=0.23 |
| Available Ca ($\mu\text{g g}^{-1}$) | 18.0 ± 14.8 | 25.9 ± 16.3 | t=0.94, p=0.36 |
| Available Mg ($\mu\text{g g}^{-1}$) | 22.1 ± 10.3 | 26.0 ± 9.9 | t=0.74, p=0.48 |

**Table S2.** Influence of wildlife (primarily pigs) on the relative abundance of the dominant bacterial families within the phyla Acidobacteria, Actinobacteria, Chlamydiae and Proteobacteria at Pasoh Forest Reserve in Peninsular Malaysia. Samples are separated by whether they were taken from open-control plots where there were many pigs versus within fenced exclosures without pigs. Only families for which significant differences were observed are shown.

| **Taxa** | **Exclosure** | **Open-Control** |
| --- | --- | --- |
| Acidobacteria |  |  |
| Solibacteraceae | 8.69 ± 1.16 | 7.11 ± 1.23 |
| Vicinamibacteraceae | 1.12 ± 0.25 | 1.89 ± 0.35 |
| Actinobacteria |  |  |
| Acidimicrobiaceae | 0.36 ± 0.16 | 0.52 ± 0.08 |
| Conexibacteraceae | 0.37 ± 0.14 | 0.58 ± 0.18 |
| Gaiellaceae | 0.11 ± 0.08 | 0.31 ± 0.11 |
| Propionibacteriaceae | 0.04 ± 0.04 | 0.15 ± 0.12 |
| Sphingobacteriaceae | 0.35 ± 0.09 | 0.58 ± 0.15 |
| Streptomycetaceae | 0.42 ± 0.25 | 1.07 ± 0.28 |
| Chlamydiae |  |  |
| Rhabdochlamydiaceae | 0.17 ± 0.07 | 0.07 ± 0.06 |
| Alphaproteobacteria |  |  |
| Bradyrhizobiaceae | 3.21 ± 0.63 | 4.74 ± 0.98 |
| Caulobacteraceae | 2.21 ± 0.86 | 4.04 ± 1.97 |
| Micropepsaceae | 1.20 ± 0.32 | 1.75 ± 0.22 |
| Rhodospirillaceae | 3.94 ± 0.90 | 5.08 ± 0.76 |
| Deltaproteobacteria |  |  |
| Polyangiaceae | 0.93 ± 0.28 | 1.73 ± 0.72 |
| Gammaproteobacteria |  |  |
| Steroidobacteraceae | 2.43 ± 0.66 | 3.34 ± 0.55 |
| Xanthomonadaceae | 0.93 ± 0.33 | 1.33 ± 0.93 |

**Figure S1:**
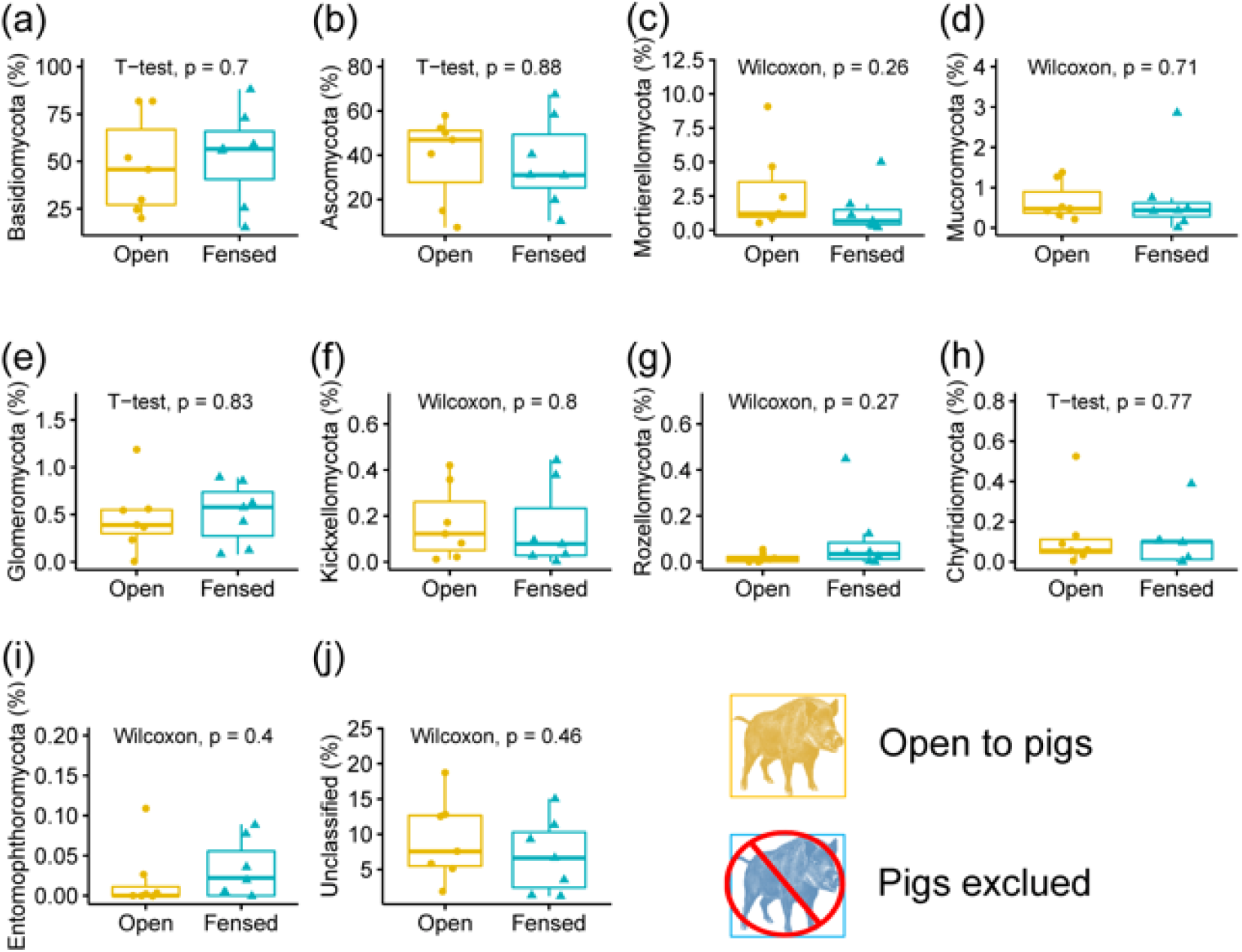
Influence of wildlife (primarily pigs) on the relative abundance of soil fungal phyla at Pasoh Forest Reserve in Peninsular Malaysia. Samples are separated by whether they were taken from open-control plots where there were many pigs (yellow dots) versus within fenced exclosures without pigs (blue triangles).

**Figure S2:**
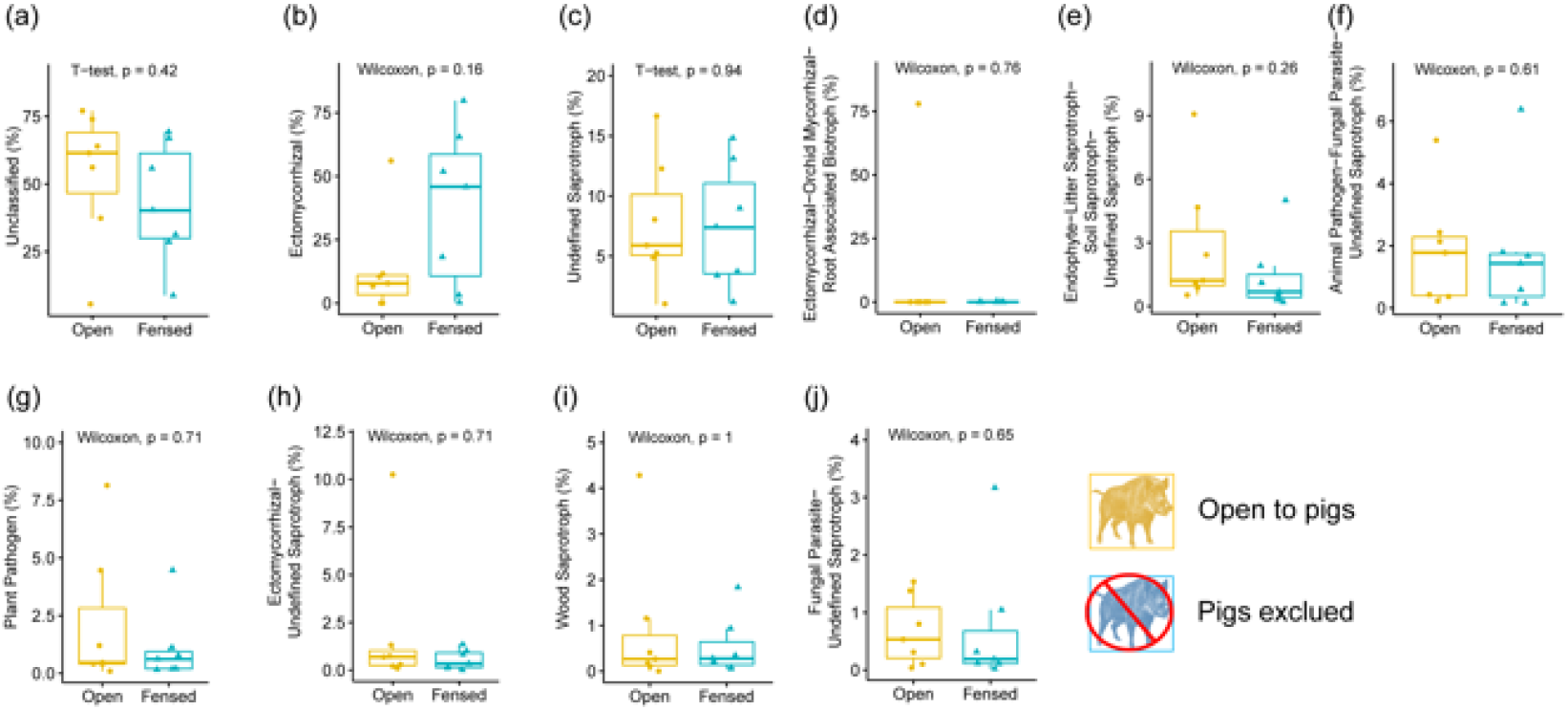
Influence of wildlife (primarily pigs) on the relative abundance of soil fungal guilds t Pasoh Forest Reserve in Peninsular Malaysia. (determined using FUNGuild) (interpretation same as Fig S1).

## Notes

https://dataview.ncbi.nlm.nih.gov/object/PRJNA748839?reviewer=ql7lqvc7ffg29i8idt4phqhfvj

### Competing Interest Statement

The authors have declared no competing interest.

